# Natural gene drives offer potential pathogen control strategies in plants

**DOI:** 10.1101/2020.04.05.026500

**Authors:** Donald M. Gardiner, Anca Rusu, Luke Barrett, Gavin C. Hunter, Kemal Kazan

## Abstract

- Globally, fungal pathogens cause enormous crop losses and current control practices are not always effective, economical or environmentally sustainable. Tools enabling genetic management of wild pathogen populations could potentially solve many problems associated with plant diseases.
- A natural gene drive from a heterologous species can be used in the globally important cereal pathogen, *Fusarium graminearum*, to remove pathogenic traits from contained populations of the fungus. The gene drive element became fixed in a freely crossing populations in only three generations.
- Repeat induce point mutation, a natural genome defence mechanism in fungi, may be useful to recall the gene drive following release, should a failsafe mechanism be required.
- We propose that gene drive technology is a potential tool to control plant pathogens.

## Introduction

Gene drives are selfish genetic elements that circumvent Mendel’s laws of independent assortment and favour their own inheritance. Because gene drives can spread rapidly through populations, they have great potential to control a variety of biological threats to plant-based agriculture, the environment and human health (Wedell *et al.*, 2019). Much of the current focus on gene drives for control of intractable pests and pathogens has concerned the molecular design of completely synthetic systems based either on nucleases such as Cas9 (DiCarlo *et al.*, 2015; Gantz *et al.*, 2015; Grunwald *et al.*, 2019) or synthetic toxin/antidote systems (Buchman *et al.*, 2018). Natural gene drives (also known as meiotic drives) have been documented in several species (Lindholm *et al.*, 2016). For fungi, the hereto cloned drive loci encode toxin-antidote type systems that protect the gametes that inherit the drive element while killing the gametes that do not inherit the drive element (Grognet *et al.*, 2014; Nuckolls *et al.*, 2017; Svedberg *et al.*, 2018; Vogan *et al.*, 2019) (Figure 1**a**). A family of related gene drive elements called spore killers (Spok) have recently been cloned from the model fungus *Podospora* spp. (Grognet *et al.*, 2014; Vogan *et al.*, 2019) with *Spok1* being the founding member.

**Figure 1:**
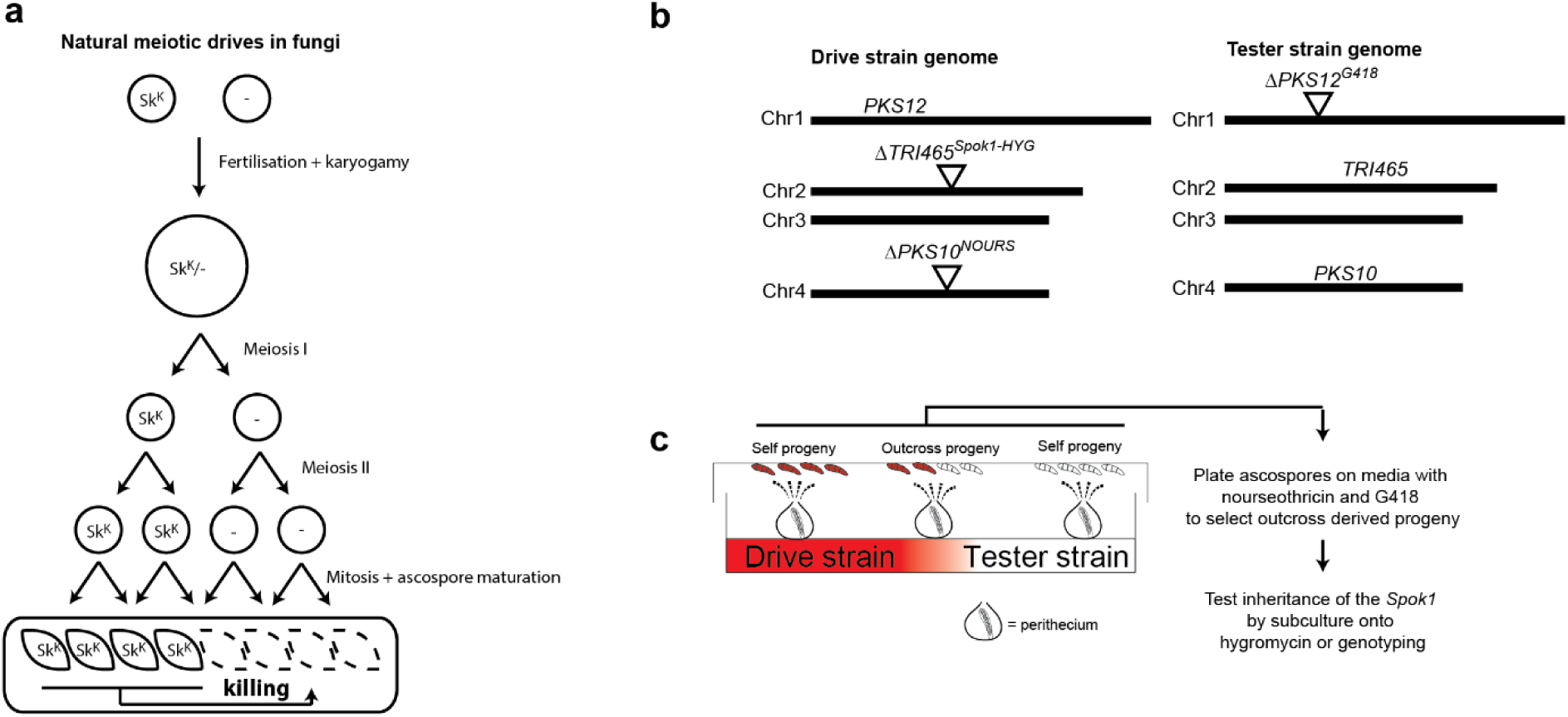
A schematic depiction of how spore killer gene drives work in fungi and the strategy used in this study to test if *Spok1* can distort the inheritance of the trichothecene toxin deoxynivalenol (DON) biosynthesis in *Fusarium graminearum*. (a) Natural gene drive systems in fungi are referred to as spore killers as they function by killing meiotic progeny who do not inherit the drive element (spore killer, Sk^K^). (b) The design of an engineered drive element in *Fusarium graminearum* where the *Spok1* spore killer gene from *Podospora comata* was used to replace three toxin genes (*TRI4, TRI5* and *TRI6* designated as *TRI465*) on chromosome 2 involved in regulation or biosynthesis of DON. Antibiotic resistance markers were placed on other chromosomes in the drive and tester strains respectively to allow isolation of recombinant progeny as *F. graminearum* can self-fertilise. (c) Crossing system for *F. graminearum* with outcrossed progeny recovered. In initial crosses, the majority of perithecia are likely to be derived from self-fertilisation. Growth of isolated ascospores on media containing nourseothricin and G418 allows for the selective isolation of progeny derived from outcrosses. *PKS10*, polyketide synthase 10 wild type locus; *ΔPKS10*^*NOURS*^, deletion mutation of *PKS10*; *TRI465*, wildtype locus for trichothecene biosynthesis genes *TRI4, TRI6* and *TRI5*; *ΔTRI465*^*Spok1-HYG*^, *Spok1* and hygromycin resistance gene in place of *TRI465* locus; *PKS12*, polyketide synthase 12 wild type locus; *ΔPKS12*^*G418*^, deletion mutation of *PKS12*.

Here, we show that such spore killer elements could be heterologously engineered to eliminate virulence traits from populations of pathogenic fungi. We selected *Fusarium graminearum* as an experimental organism, as this globally important cereal pathogen causing Fusarium head blight (FHB) disease (Goswami & Kistler, 2004) can reproduce sexually and many genes associated with virulence have been characterized (Kazan *et al.*, 2012). As a well-known virulence factor, the trichothecene (TRI) mycotoxin deoxynivalenol (DON) and its derivatives, produced by *F. graminearum* during infection, contribute to disease development on wheat, significantly reducing grain quality and causing adverse health effects in humans and animals who consume contaminated grains (Goswami & Kistler, 2004). We show that *Spok1* can be used to distort inheritance at two genomic loci (the *TRI* gene cluster and *ABC1*) that control virulence in *F. graminearum*. As *F. graminearum* can undergo both selfing and outcrossing, we show that even with both modes of sexual reproduction occurring simultaneously in a contained population, the gene drive element can reach fixation in only a few generations. We also demonstrated that repeat induced point mutation (RIP), a fungal defence mechanism, could be harnessed to limit the functionality of the drive if required.

## Materials and methods

### Strain generation

All transformants were created in the *F. graminearum* isolate CS3005 background (Gardiner *et al.*, 2014). Transformation was performed as previously described (Desmond *et al.*, 2008). Vectors were all designed for double cross-over replacement of the target locus with full details in the supplementary materials. The strains created and used are summarised in Table 1. All transformants were initially screened using triplex PCR assays (primers in Supplementary Table 1) and in selected transformants whole genome sequenced to verify deletion of the target genes and single copy status of the inserted DNA as described previously (Gardiner & Kazan, 2018). Sequence reads for the transformants have been deposited in the GenBank sequence read archive under BioProjects PRJNA591033 and PRJNA605650. The wild type and mutant loci along with regions of sequence homology used for double crossover replacements and primer binding sites for mutant screening, along with read mapping graphs for one selected transformant, for all four vectors are presented in Supplementary Figure 1. For all constructs, except for the *ABC1* replacement, two independent single copy mutant strains were used in experiments but no differences between strains with the same genotype were observed and as such the nomenclature used herein refers to the genotype and not transformant number.

**Table 1:** Strains used in this study

| Nomenclature | Description | Reference |
| --- | --- | --- |
| $\Delta TRI465^{Spok1-HYG}$ | Transformants with *TRI4*, *TRI6* and *TRI5* replaced with a 3.5 Kbp genomic fragment containing *Spok1* and also a hygromycin (HYG) resistance gene driven by the *Aspergillus nidulans* *TrpC* promoter | This study |
| $\Delta PKS10^{NOURS}$ | Deletion mutant of polyketide synthase 10 with a *A. nidulans* β-tubulin promoter driven nourseothricin (NOURS) resistance gene | This study |
| $\Delta PKS12^{G418}$ | Deletion mutant of polyketide synthase 12 with a *A. nidulans poly-ubiquitin* promoter driven nourseothricin (NOURS) resistance gene | This study |
| $\Delta ABC1^{Spok1-oHYG}$ | A transformant where the ABC1 locus was replaced with a 3.5 Kbp genomic fragment containing *Spok1* and also a codon optimised hygromycin (HYG) resistance gene driven by the *A. nidulans cyp51* promoter | This study |
| $\Delta TRI465^{Cas9sgRNA::TRI5}$ | Transformants with *TRI4*, *TRI6* and *TRI5* replaced with a *Cas9* and synthetic guide RNA targeting *TRI5* plus a hygromycin resistance gene | (Gardiner & Kazan, 2018) |

### Sexual crosses

Crossing was carried out as previously described (Cavinder *et al.*, 2012) except that half-strength carrot agar was used (130 g peeled carrots with deionised water to 1 L, blended to a smooth consistency in a Breville The Boss blender for 2 minutes on high with 15 g L^-1^ agar, autoclaved once). Ascospores were harvested by washing from Petri dish lids and plated on 10% PDA amended as appropriate with antibiotics. Drive strains were created by crossing the *ΔPKS10*^*Tub-NOURS*^ and *ΔTRI465*^*Spok1-TrpC-HYG*^ or *ΔABC1*^*Spok1-oHYG*^ transformants with selection on 50 mg L^-1^ of nourseothricin and 200 mg L^-1^ hygromycin. For assessing drive from outcrossing, ascospores were initially plated on G418 and nourseothricin (both at 50 mg L^-1^). No antibiotics were used for assessing drive performance in crosses undergoing random mating. Ascospores were germinated overnight in a 23°C incubator (Memmert, Schwabach, Germany) and individuals were picked using an Olympus (Tokyo, Japan) SZ51 stereo microscope onto separate Petri plates containing 10% PDA. After additional subculture onto solid media containing either G418, nourseothricin or hygromycin, the presence of absence of growth from these individual ascospore derived colonies, on the respective antibiotics, was used to infer the genotype for each of the three loci used in the crosses.

### Analysis of repeat induced point mutation

To test if RIP could be used as a tool to mutate the *Spok1* sequence, a strain was constructed via crossing to bring together the two insertions of *Spok1*, one at the *TRI465* locus and the other at the *ABC1* locus. Progenies were screened using the same sets of primers used to initially screen the knockout strains. The strain containing both insertions of *Spok1* was subcultured twice on half-strength carrot agar and ascospores harvested from the lid of the second subculture. A 1200 bp portion of the two *Spok1* copies were initially amplified together and sequenced (primers DG1330 and DG1333). Subsequently the two copies of *Spok1* from a single progeny were amplified and sequenced independently. Amplification of the insertion of *Spok1* in the *ABC1* locus occurred using primers DG1324 and DG1207. Amplification of the insertion of *Spok1* in the *TRI465* locus used DG1178 and DG1217. Phusion DNA polymerase with annealing at 63°C was used according the manufacturer’s instructions (NEB). Both products were sequenced with DG1328-DG1333 using Sanger sequencing performed by the Australian Genome Research Facility.

### Statistical analyses

Segregation was analysed using Goodness of Fit test (Chi-Square) in Microsoft Excel. The Chi-square statistics were calculated using custom formulae and *p*-values determined using the CHISQ.TEST function for observed versus expected frequencies.

## Results

### *Spok1* is functional when inserted the trichothecene cluster in *F. graminearum*

We first established a system to test the ability of the *Spok1* to act as a heterologous meiotic distorter in *F. graminearum* by replacing three adjacent genes (*TRI4, TRI5* and *TRI6*) of the *TRI* gene cluster required for DON production with *Spok1* (Figure 1**b**). As *F. graminearum* can both self-fertilise and outcross, the test system was designed to distinguish progeny derived from these two modes of sexual reproduction. Of twenty-six recombinants identified from a cross between the drive (*Spok1* present) and tester (*Spok1* absent) strains, twenty-four inherited the *ΔTRI465*^*Spok1-HYG*^ allele, suggesting non-equal segregation (*p*-value 2×10^−5^) with a drive efficiency of ∼92%. This finding shows that Spok1 is functional in *F. graminearum*. When a strain expressing *Cas9* and a *TRI5* targeting guide RNA (*ΔTRI465*^*Cas9-sgRNA::TRI5*^) was used in place of the *Spok1* insert in the *TRI465* locus (Gardiner & Kazan, 2018) equal segregation (8:8) was observed strongly suggesting this particular nuclease based design, similar to high efficiency drive systems shown to work in a number of other species including yeast and insects (DiCarlo *et al.*, 2015; Hammond *et al.*, 2016), is unable to drive in *F. graminearum*.

### *Spok1* also functions in the *ABC1* virulence locus in *F. graminearum*

In *Podospora spp.*, the proximity to the centromere affects the drive efficiency of *Spok1* with positioning closer to the centromere resulting in stronger drive (Grognet *et al.*, 2014). This may be due to intrinsic features of the final stages of ascospore generation in *Podospora spp*. which results in binucleated spores. To test if drive could be achieved when placed at an alternative genomic locus in *F. graminearum*, a strain was created whereby *Spok1* replaced the *ABC1* gene encoding an ABC (ATP-binding cassette) transporter shown to have a role in virulence and xenobiotic tolerance (Abou Ammar *et al.*, 2013; Gardiner *et al.*, 2013). The *ABC1* gene is located approximately 0.5 Mbp from the end of chromosome 2. This position is ∼5.2 Mbp (∼347 cM) from the centromere of this chromosome (King *et al.*, 2015; Laurent *et al.*, 2018). In contrast, the *TRI465* locus is ∼2.1 Mbp (∼195 cM) from the centromere but on the same arm of this chromosome as *ABC1* (King *et al.*, 2015; Laurent *et al.*, 2018). In crosses selecting only recombinant progeny, 45/54 progeny inherited the *Spok1* locus, suggesting a drive efficiency of 83% which was significantly different to that expected under neutral segregation (*p*-value 9×10^−7^). The difference in drive efficiency observed between the *TRI465* and *ABC1* locus was significant (*p*-value 0.01) although the biological reason for this is unclear, as unlike the case in *Podospora*, both loci are essentially genetically unlinked from the centromere.

### Selfing is not necessarily an impediment to drive spread

*Fusarium graminearum* is a homothallic fungus, meaning that it undergoes selfing and outcrossing. In nature, both modes of reproduction are known to occur (Goswami & Kistler, 2004; Trail, 2009), and while outcrossing promotes spread, selfing could potentially limit the drive’s spread in populations (Bull *et al.*, 2019). To test if drive could also be observed under random mating in *F. graminearum*, we crossed the drive and tester strains and monitored resulting genotypes in randomly mating populations (i.e. without any antibiotic selection) over three successive generations (Figure 2**a**). In the first generation of this cross, 16 out of 50 recovered progeny had recombinant genotypes (Figure 2**b**). Of these 16, 15 inherited the *ΔTRI465*^*Spok1-HYG*^ allele, corresponding to a drive efficiency of ∼94%. Although the selfing rate could not be directly measured in this experimental setup after the first generation, in the second generation, 52% of strains had genotypes that differed from the original parental strains, suggesting continued outcrossing in the population. In the second generation, all strains that had non-parental genotypes carried *ΔTRI465*^*Spok1-HYG*^. By the third generation none of the sampled strains contained the tester parent genotype. Rather, *ΔTRI465*^*Spok1-HYG*^ was fixed in the population (n=50) while the other non-driving engineered loci (*ΔPKS10*^*NOURS*^ and *ΔPKS12*^*G418*^) were observed in 44% and 54% of the progeny, respectively (Figure 2**b**). In a separate experiment, using an independent *ΔTRI465*^*Spok1-HYG*^ transformant, despite initially only recovering just 6% recombinant progeny, a similar trend was observed with *ΔTRI465*^*Spok1-HYG*^ present in 82% of all progeny by the third generation (Supplementary Figure 2). Thus, even in the presence of a mixed-mating systems, the drive was able to spread, although in this *in vitro* scenario, there is no fitness cost associated with the drive’s cargo (i.e. inability to produce toxin).

**Figure 2:**
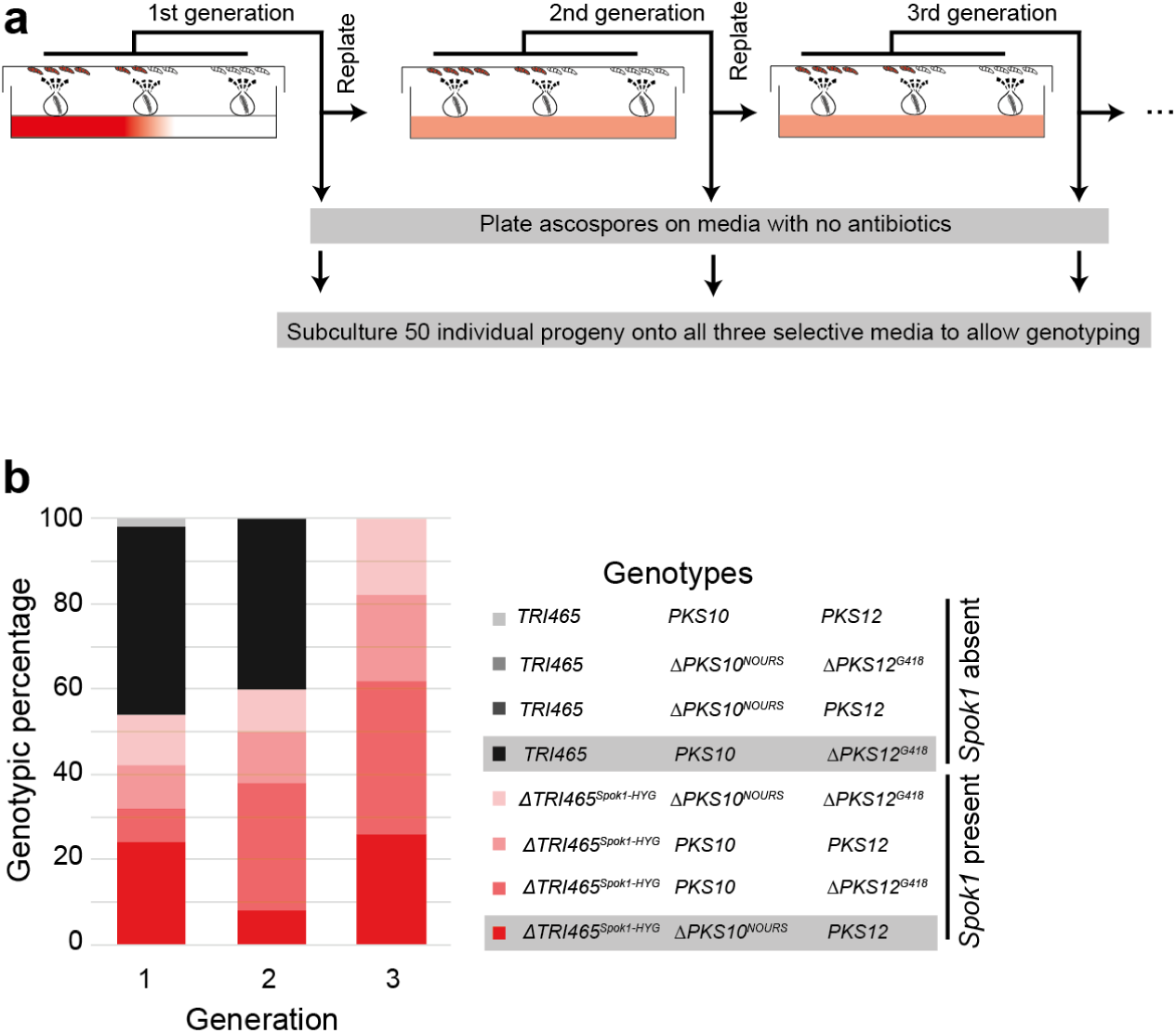
Multi-generation analysis of outcrossing and *Spok1* spread in *Fusarium graminearum*. (a) The system to follow the inheritance of the *Spok1* element in a randomly mating population where both self-fertilisation and outcrossing are possible. The system was established with a cross between the drive and tester strains with progeny monitored across three generations. At each generation, progenies were harvested, and the majority of the population was replated directly onto carrot agar to initiate the next cycle. At each generation, a small subsample of the population was germinated on media lacking antibiotics and 50 germinated ascospores were picked and subsequently phenotyped for resistance to hygromycin (indicative of carrying *Spok1*), nourseothricin (indicative of carrying ΔPKS10) and G418 (indicative of carrying ΔPKS12). (b) Genotypic analysis of the 50 ascospores sampled at each generation. With three loci segregating in these populations, in theory eight possible genotypes are possible. The genotypes of the two parental strains are shaded in grey in the legend. *PKS10*, polyketide synthase 10 wild type locus; *ΔPKS10*^*NOURS*^, deletion mutation of *PKS10*; *TRI465*, wildtype locus for trichothecene biosynthesis genes *TRI4, TRI6* and *TRI5*; *ΔTRI465*^*Spok1-HYG*^, S*pok1* and hygromycin resistance gene in place of *TRI465* locus; *PKS12*, polyketide synthase 12 wild type locus; *ΔPKS12*^*G418*^, deletion mutation of *PKS12*.

### The natural genome defence, repeat induced point mutation, could be used to recall the gene drive

The possible application of gene drives must be tempered by careful consideration of any risks that may result from their use and groups working on synthetic drives have called for highly cautious approaches both in terms of global regulation and molecular design (Esvelt & Gemmell, 2017). Therefore, tools that enable recall of the gene drive after environmental release will likely be important components of safe drive systems and help achieve social and regulatory acceptance of gene-drive technologies. One suggested solution to this issue is the release of a second gene drive, engineered to render the original drive non-functional. Modelling suggests that in this scenario, the fitness cost of the original drive needs to be high while the kill drive’s effect on fitness needs to be negligible (Girardin *et al.*, 2019). We hypothesised that repeat induced point mutation (RIP), a natural genome defence mechanism operating in many sexual fungi, could be harnessed for this purpose. RIP causes C→T mutations in sequences that contain repeats greater than ∼400 bp (Gladyshev, 2017) located anywhere in the genome. The recognition and mutation of repetitive sequences within the parental nucleus occurs following cell fusion but prior to nuclear fusion during meiosis (Gladyshev, 2017). RIP is moderately active in *F. graminearum* (Pomraning *et al.*, 2013) and likely to have been a key process in shaping the repeat-poor genome of this species (Cuomo *et al.*, 2007).

To test the utility of RIP as a drive recall option, we constructed another strain by crossing the two drive strains that carry two separately engineered *Spok1* sequences in place of both the *TRI465* and *ABC1* loci. In this strain, RIP mutations would be expected to occur in subsequent sexual cycles (selfing or outcrossing) until the homology between the two repeated *Spok1* sequences drops below 80% (Figure 3**a**). Ascospores derived from strains carrying *Spok1* at both loci were initially analysed by amplification of *Spok1* sequences from both loci and then sequenced by Sanger technology. The occurrence of mutations was identified in multiple strains by the presence of chromatograms with mixed nucleotide calls identified by manual inspection. Subsequently, the two *Spok1* copies from a single progeny were amplified independently and the 2206 bp *Spok1* coding region was sequenced. As shown in Figure 3**b**, amino acid changing mutations were observed in both copies from this strain. Whether these amino acid changes render the drive element non-functional are not known. However, the P620S change observed in the Spok1 copy from the *TRI465* locus occurred in a residue previously identified as being highly conserved in Spok family members (Vogan *et al.*, 2019). While the average number of generations required to remove the functionality of Spok1 needs to be determined, our data demonstrate RIP is capable of rapidly inducing changes in coding sequence of both copies of the drive elements. Therefore, even the moderately active nature of RIP in *F. graminearum* could provide a system to render Spok1 function to be slowly lost from a population over several generations.

**Figure 3.**
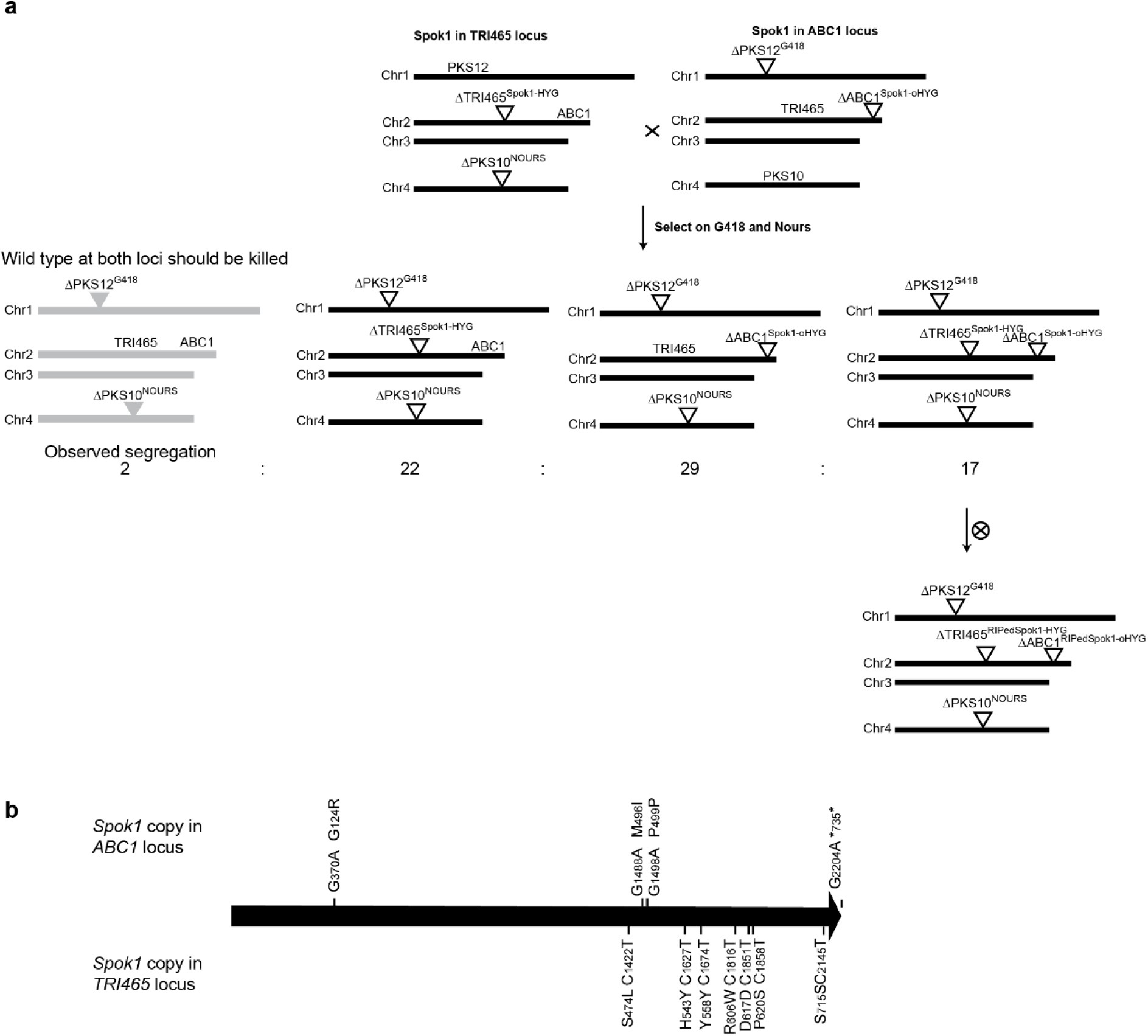
(a) A strategy to recall the drive element by exploiting repeat induced point mutation (RIP), a natural genome defence mechanism in fungi. This strategy is based on the expectation that in fungal strains in which the *Spok1* gene is repeated in the genome, subsequent sexual cycles should render the repeated sequence inactive. Here, to bring together *Spok1* in a new strain, ΔTRI465^Spok1-HYG^ and *ΔABC1*^*Spok1-oHYG*^ strains were crossed and the progeny selected for G418 and NOURS resistance. The hygromycin phospho-transferase alleles used in parental strains differ and should not be mutated by this process. Progeny from this cross segregated in 2:22:29:17 ratios for the four possible genotypes (TRI465^WT^, ABC1^WT^:ΔTRI465^Spok1-HYG^, ABC1^WT^:TRI465^WT^, ΔABC1^Spok1-oHYG^:ΔTRI465^Spok1-HYG^, ΔABC1^Spok1-oHYG^ (b) The *Spok1* copies present in a single ascospore derived strain from a self-fertilisation of a strain carrying *Spok1* at both the *ABC1* and *TRI465* loci were sequenced. The *Spok1* copy at the *ABC1* locus contained three nucleotide changes and the *Spok1* copy from the *TRI465* locus contained six changes. The nucleotide changes and resulting coding consequences are shown above (*ABC1* copy) and below (*TRI465* copy) the arrow representing the *Spok1* coding sequence.

## Discussion

In this study, we showed that *Spok1*, a natural gene drive locus previously identified in the model fungus *Podospora*, could successfully distort the inheritance of two separate virulence-associated loci in the pathogenic fungus *F. graminearum*, despite these two fungi sharing a last common ancestor c130 Mya (Prieto & Wedin, 2013). Given this finding, Spok1 may also function in a suite of other fungal pathogens. With regard to *F. graminearum*, testing the efficacy of this system in the field is currently not possible due to social and regulatory constraints. However, considerable experimental and pathogen-specific modelling studies can be undertaken to predict how this approach would work under field conditions. Certainly, *F. graminearum* can spread relatively quickly in agricultural settings. This pathogen appears to have originated in North America, presumably associated with a native grass (Lofgren *et al.*, 2018). Over the last 400 years, since wheat was first brought to North America (Lofgren *et al.*, 2018), *F. graminearum* has spread globally. On even shorter time-scales, more toxigenic lineages of *F. graminearum* seem to be displacing others (Ward *et al.*, 2008) meaning drive-containing strains may have the potential to spread relatively rapidly from initial deployment, under a reasonable rate of outcrossing in the field. As indicated above, *F. graminearum* selected as a model pathogen to demonstrate the utility of this technology, is a homothallic fungus. This strategy might work more efficiently in strictly heterothallic fungi that undergo extensive sexual recombination in nature such as *Leptosphaeria maculans*, the causative agent of the blackleg disease in oilseed rape (Rouxel & Balesdent, 2005), or *Zymoseptoria tritici* the causal agent of Septoria blotch of wheat (Orton *et al.*, 2011).

One of the virulence-related loci we targeted here was DON production, which causes significant health issues when ingested by humans and animals. Therefore, elimination of DON and other mycotoxins from food and feed has long been a major objective of researchers, food producers, regulators and public health experts. Current mechanisms for controlling dietary exposure to mycotoxins are cumbersome and multilayered (Cheli *et al.*, 2017) and this work demonstrates that gene drive strategies are worthy of consideration to manage toxin contamination issues. Nevertheless, further understanding of the ecological role that toxin production by *F. graminearum* plays in nature may be required.

Although we have demonstrated that natural meiotic distorter loci have the potential to eliminate unwanted genes from contained populations, a drive such as this is also capable of delivery of “cargo genes” or “genetic cargo” into fungal populations. Indeed, the antibiotic resistance cassette used here to integrate the *Spok1* gene into the pathogen genome is technically a cargo gene. Other examples of cargo genes that are relevant to agriculture and plant protection could be pathogen avirulence genes or fungicide sensitivity genes/alleles in species where these traits are relevant. Spreading avirulence genes into pathogen populations would enable their detection by corresponding resistance genes already deployed in crops, or present in wild plant populations (Barrett *et al.*, 2019). Similarly, making pathogen populations re-sensitised to existing fungicides could renew abandoned chemical control options. However, these measures exert strong selection pressure on pathogen populations and may lead to changes to overcome the effect of such measures. Quantitative reduction in virulence may prove to be a more sustainable approach in the long term.

In conclusion, this study has demonstrated that a natural spore killer element can function in a distantly related pathogenic fungus by distorting the inheritance of virulence loci. In addition, we show that natural genome defence mechanisms could be harnessed to render the gene drive non-functional if required. Genetic elements like this could form the basis of future, sustainable agricultural systems with a decreased reliance of chemical control measures. It is envisaged that reducing virulence would minimize crop losses without driving pathogen populations into extinction, a concern currently associated with the use of gene drives. However, additional work is needed to determine the mode of action of spore killers and/or other similar meiotic drives for scientific as well as regulatory reasons. Although the technology undoubtedly involves the release of a genetically modified organism, the potentially transformative impact of this type of technology, should at least be investigated further with appropriate regulation, alongside studies to gauge social acceptance and safety of such technologies (Jones *et al.*, 2019).

## Supplementary materials

### Strain generation

The construct for replacing *TRI465* (*FG05_03535, FG05_16251 and FG05_03537*), which resides on chromosome 2, with *Spok1-HYG* was generated by assembling fragments using yeast recombinatorial cloning. One of the flanks for homologous recombination was 1389 bp of *F. graminearum* isolate CS3005 sequence corresponding to the *TRI4* terminator region and was amplified using DG1168 and DG1239. The *Spok1* genomic fragment, corresponding to 3540 bases of accession JX560967 (nucleotide position 4744-8283), was synthesised flanked by *Hpa*I/*Sal*I sites by Genscript and released with a *Hpa*I/*Sal*I digest. The *TrpC-Hygromycin phosphotransferase* sequence and 957 bp of the region downstream of the *TRI5* stop codon (used as the other flank for homologous integration) was amplified using DG1240 and DG1167 in a single piece from a plasmid corresponding to GenBank accession MF084286.1. Fragments were transformed into yeast and subsequently *F. graminearum* as described above. The genotype of the strain carrying *Spok1* at the *TRI465* locus is termed *ΔTRI465*^*Spok1-HYG*^ throughout the manuscript. An equivalent insertion with Cas9 and a guide RNA targeting *TRI5* was previously described (Gardiner & Kazan, 2018).

We also engineered a strain to carry a nourseothricin (NOURS) resistance cassette on chromosome 4 in place of *polyketide synthase 10* (*PKS10*) responsible for fusarin production (Brown *et al.*, 2012) to facilitate the selection of ascospores resulting from outcrossing. Knockout mutants of *PKS10* show normal pathogenicity on wheat (Gaffoor *et al.*, 2005). The vector for creating the *ΔPKS10*^*NOURS*^ strain was synthesised by Genscript (USA). The vector consisted of 1000 bp of sequence upstream of the *PKS10* (*FG05_07798*) coding sequence, 800 bp of the *Aspergillus nidulans beta-tubulin* (*ANIA_01182*) promoter derived from the genome sequence for this organism (Wortman *et al.*, 2009), the *nourseothricin acetyl transferase* gene from *Streptomyces nousei* and 1000 bp of sequence downstream of the stop codon of *PKS10*. The *beta-tubulin-nourseothricin acetyl transferase* cassette has been submitted to GenBank under accession MK431404. The genotype of strains carrying this deletion at the *PKS10* locus is represented as *ΔPKS10*^*NOURS*^.

In the tester strain (Figure 1), we replaced the gene encoding the red pigment polyketide synthase (*PKS12*) (Malz *et al.*, 2005) on chromosome 1 with a geneticin (G418) resistance marker, which rendered the strain albino. Knockout mutants of *PKS12* show normal pathogenicity on cereals (Gaffoor *et al.*, 2005; Malz *et al.*, 2005). The vector for creating the *ΔPKS12*^*G418*^ strain was generated by assembling PCR-derived fragments using yeast recombinatorial cloning. The 5’ flank of *PKS12* (*FG05_02324*) was amplified with DG1174 and DG1184 from gDNA of CS3005. The *A. nidulans poly-ubiquitin* promoter (*ANIA_02000*) was amplified from a fragment synthesised by Genscript with DG1185 and DG1186 to give a 1313 bp regulatory region driving the *neomycin phosphotransferase* gene (*NPTII*) which was amplified along with 900 bp of sequence downstream of *PKS12* using DG1187 and DG1177 from a synthesised plasmid. All fragments, with a *Hin*dIII/*Xba*I cut preparation of pYES2, were transformed into yeast strain BY4743 as described elsewhere (Gietz & Schiestl, 2007). A maxi scale preparation (QIAgen HiSpeed maxi prep kit) was used for fungal transformation. The *poly ubiquitin-NPTII* cassette has been submitted to GenBank under accession MK431401. The genotype of strains carrying this deletion at the *PKS12* locus is represented as *ΔPKS12*^*G418*^.

The vector for replacing *ABC1* with *Spok1* was generated by assembling fragments using yeast recombinatorial cloning. A region corresponding to 1464 to 190 bp upstream of the *ABC1* start codon was amplified using primers DG1314 andDG1315 and used as the 5’ homology flank for the knockout generation. A 595 bp *A. nidulans cytochrome P450 monoogygenase 51B* (*ANIA_08283*) promoter and *F. graminearum* codon optimised *hygromycin phosphotransferase* (*oHPT*) encoding gene were originally obtained as a gBLOCK (IDT, Singapore) and subsequently amplified with DG1316 and DG1317. The sequence of this cassette has been deposited in GenBank (MK431402). The *Spok1* fragment was identical to that used in the *TRI465* replacement above. The 3’ flank of *ABC1* was amplified with DG1318 and DG1319 corresponding to 1084 bp immediately downstream of the *ABC1* stop codon. A linearized (*Xba*I) maxi-scale preparation (QIAgen HiSpeed maxi prep kit) was used for fungal transformation. The genotype of strains carrying *Spok1* in the ABC1 locus is represented as *ΔABC1Spok1oHYG*.

## Supplementary information

**Supplementary Figure 1:**
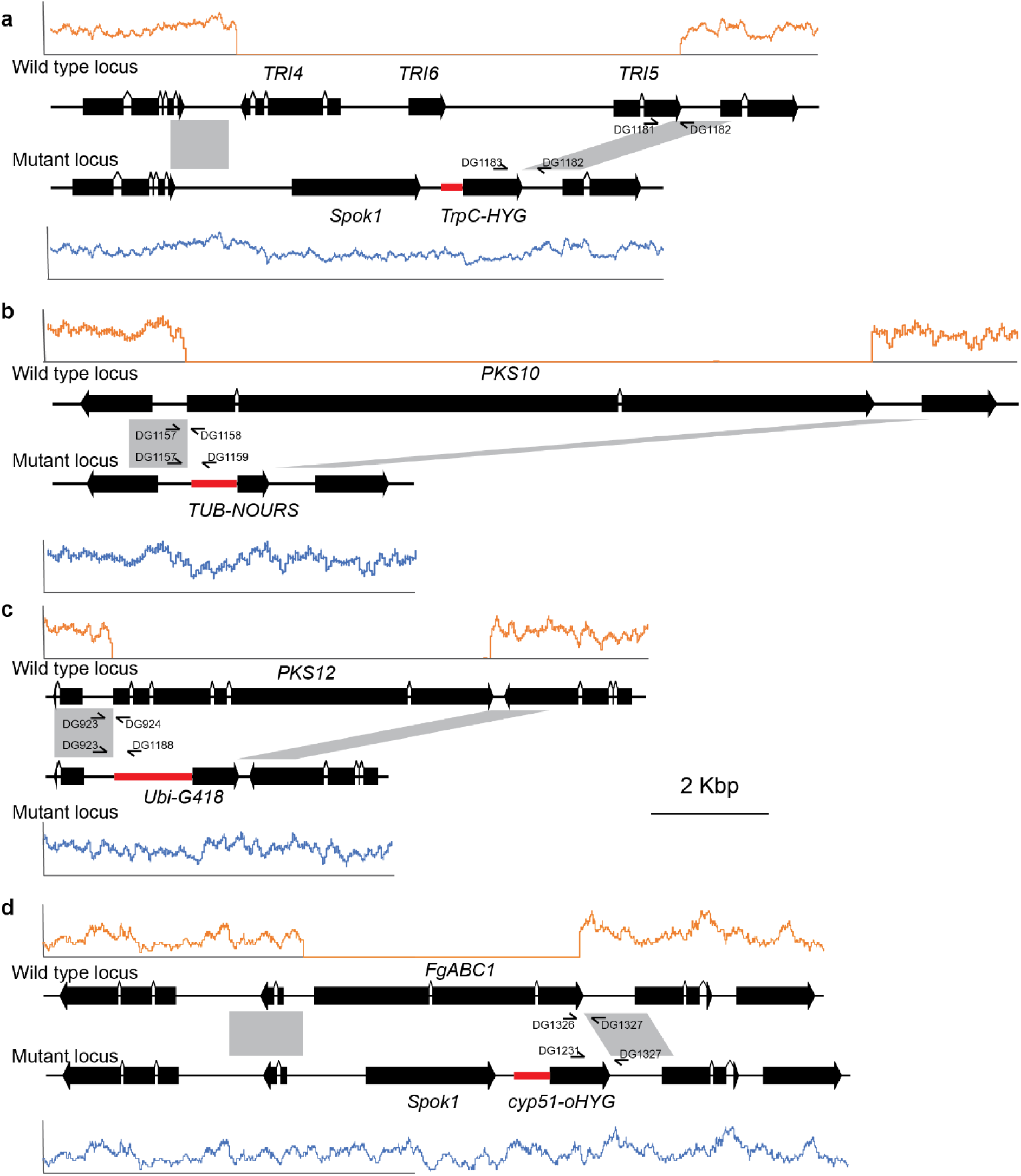
Wild type and mutant loci for the five loci targeted by homologous integration in this work. For each loci, reads from the mutant strains were mapped to the wild type genome and expected transformant genome to determine correct insertion of the transgene. Shown are read coverage graphs for one selected mutant mapped to the CS3005 parental locus (in orange above the locus diagram) and the expected transformant locus (in blue below the locus diagram). Promoters from *Aspergillus nidulans* are coloured red. Grey boxes between the wild type and mutant loci represent the regions used in the vectors for homologous integration at the target site. Primers binding sites used for screening of mutants are represented below the wild type loci and above the mutant loci. (a) The insertion of *Spok1* at the *TRI465* locus using the TrpC-hygromycin phosphotransferase (designated as *TrpC-HYG*) to select for transformants. (b) Replacement of PKS10 with the tubulin promoter (TUB)-nourseothricin acetyl transferase cassette (designated as *TUB-NOURS*). (c) Replacement of the *PKS12* gene with the poly ubiquitin promoter (*Ubi)-neomycin phosphotransferase* cassette (designated as *Ubi-G418*). (D) Replacement of the *ABC1* gene with *Spok1* and the *cytochrome P450 monooxygenase 51* promoter (*cyp51*)-codon optimised hygromycin phosphotransferase cassette (designated as *cyp51-oHYG*).

**Supplementary Figure 2:**
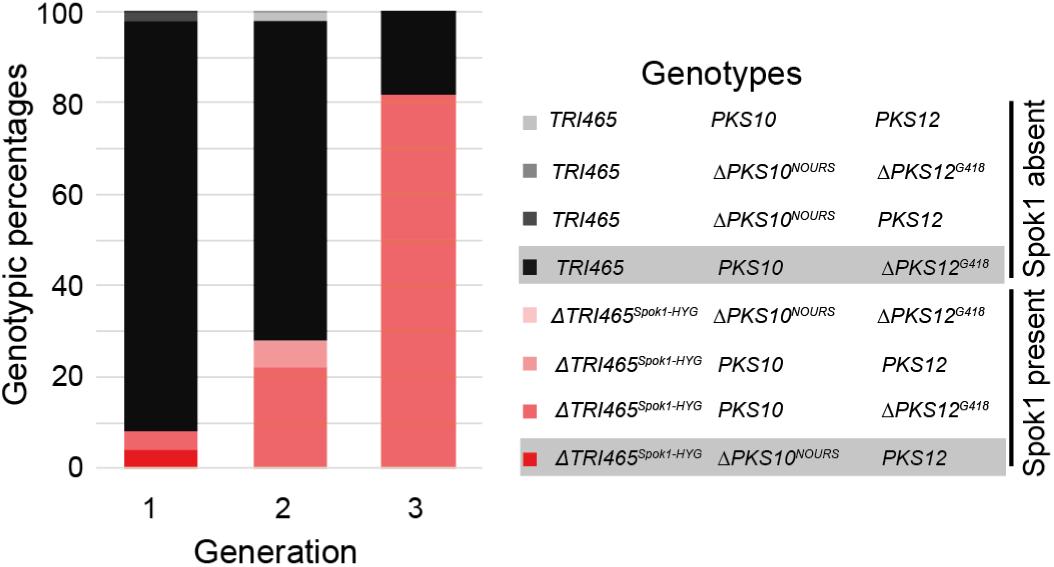
Repeated experiment of a multi-generation analysis of outcrossing and *Spok1* spread in *Fusarium graminearum*. A randomly mating population was established between the drive and tester strains and monitored across three generations. At each generation, progenies were harvested and germinated on media lacking antibiotics. Fifty germinated ascospores were picked at each generation and the resultant cultures subsequently phenotyped for resistance to hygromycin (indicative of carrying *Spok1*), nourseothricin (indicative of carrying ΔPKS10) and G418 (indicative of carrying ΔPKS12). The genotypes of the two parental strains are shaded in grey in the legend.

**Supplementary Table 1:**
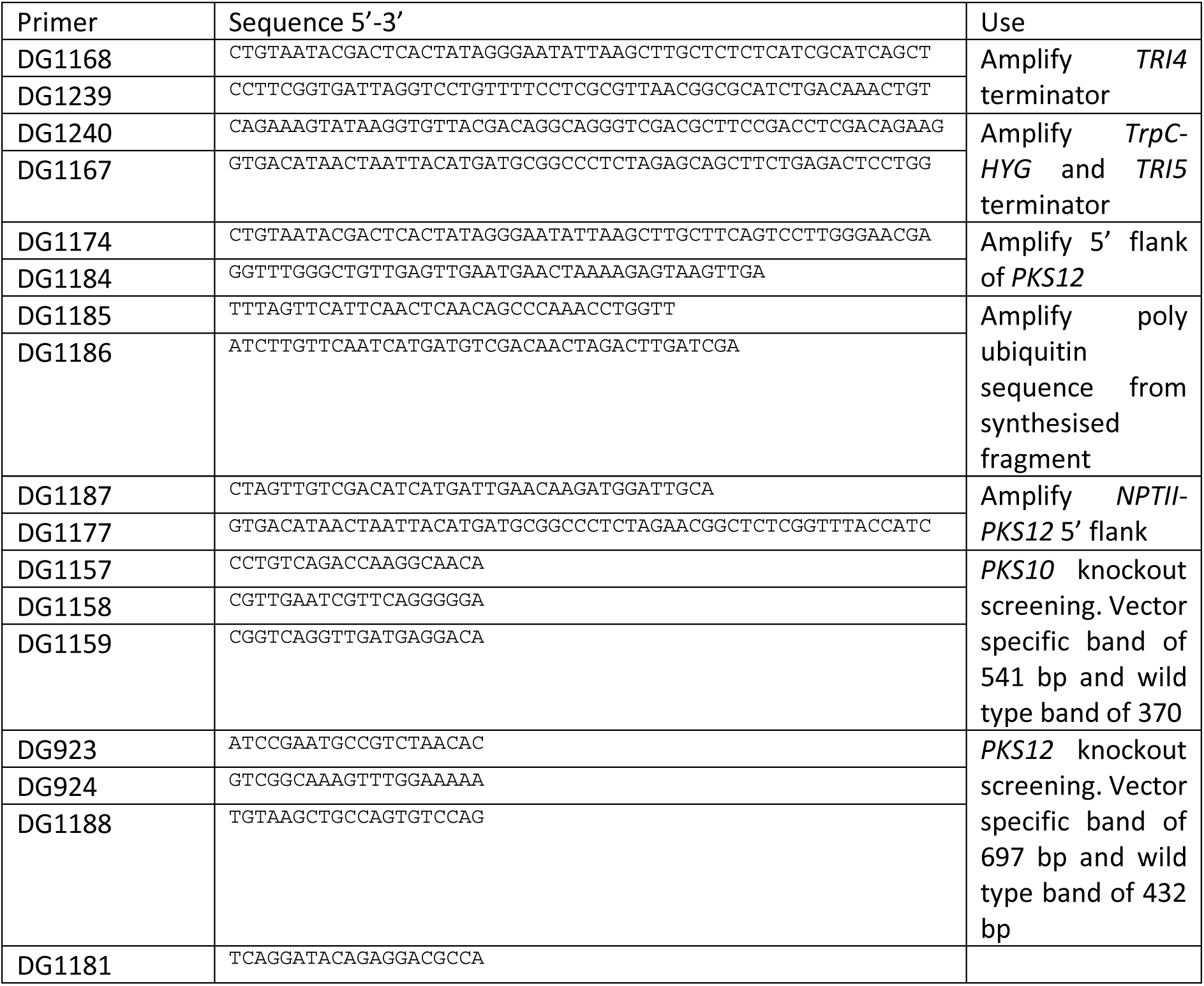
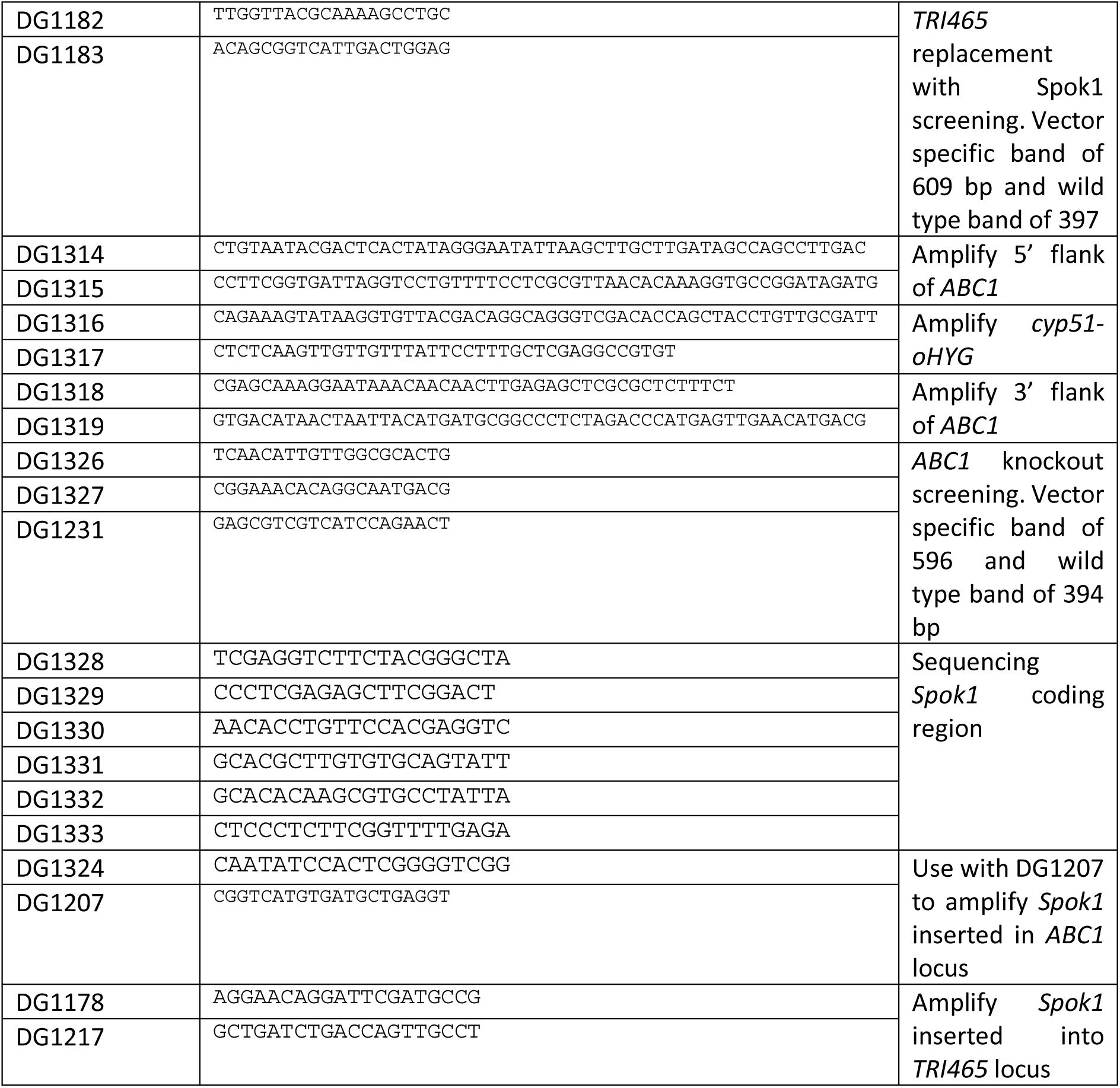
Primers used for construction of plasmids and screening of transformants for correct integration.

## Acknowledgements and funding

This work was solely funded by CSIRO. The authors wish to thank Prof. Michael Freitag (Oregon State University) for useful discussions around repeat induced point mutation.

## Author contribution

DMG, LB, GCH, KK planned and designed the research. DMG, AR performed experiments and analysed data. DMG, AR, LB, GCH, KK wrote the manuscript.

## References

Abou Ammar G, Tryono R, Döll K, Karlovsky P, Deising HB, Wirsel SGR. 2013. Identification of ABC transporter genes of *Fusarium graminearum* with roles in azole tolerance and/or virulence. PLoS ONE 8(11): e79042.

Barrett LG, Legros M, Kumaran N, Glassop D, Raghu S, Gardiner DM. 2019. Gene drives in plants: opportunities and challenges for weed control and engineered resilience. Proceedings of the Royal Society B: Biological Sciences 286(1911): 20191515.

Brown DW, Butchko RAE, Busman M, Proctor RH. 2012. Identification of gene clusters associated with fusaric acid, fusarin, and perithecial pigment production In *Fusarium verticillioides*. Fungal Genetics and Biology 49(7): 521–532.

Buchman A, Marshall JM, Ostrovski D, Yang T, Akbari OS. 2018. Synthetically engineered *Medea* gene drive system in the worldwide crop pest *Drosophila suzukii*. Proceedings of the National Academy of Sciences 115(18): 4725–4730.

Bull JJ, Remien CH, Krone SM. 2019. Gene-drive-mediated extinction is thwarted by population structure and evolution of sib mating. Evolution, Medicine, and Public Health 2019(1): 66–81.

Cavinder B, Sikhakolli U, Fellows KM, Trail F. 2012. Sexual development and ascospore discharge In Fusarium graminearum. Journal of Visual Experiments 61: e3895.

Cheli F, Pinotti L, Novacco M, Ottoboni M, Tretola M, Dell’Orto V 2017. Mycotoxins in wheat and mitigation measures. In: Wanyera R, Owuache J eds. Wheat improvement, management and utilization. London: IntechOpen Limited, 227–251.

Cuomo CA, Güldener U, Xu JR, Trail F, Turgeon BG, Di Pietro A, Walton JD, Ma LJ, Baker SE, Rep M, et al. 2007. The *Fusarium graminearum* genome reveals a link between localized polymorphism and pathogen specialization. Science 317.

Desmond OJ, Manners JM, Stephens AE, Maclean DJ, Schenk PM, Gardiner DM, Munn AL, Kazan K. 2008. The *Fusarium* mycotoxin deoxynivalenol elicits hydrogen peroxide production, programmed cell death and defence responses in wheat. Molecular Plant Pathology 9(4): 435–445.

DiCarlo JE, Chavez A, Dietz SL, Esvelt KM, Church GM. 2015. Safeguarding CRISPR-Cas9 gene drives in yeast. Nat Biotech 33(12): 1250–1255.

Esvelt KM, Gemmell NJ. 2017. Conservation demands safe gene drive. PLoS Biology 15(11): e2003850.

Gaffoor I, Brown DW, Trail F. 2005. Functional analysis of the polyketide synthase genes in the filamentous fungus *Gibberella zeae* (anamorph *Fusarium graminearum*). Eukaryotic Cell 4.

Gantz VM, Jasinskiene N, Tatarenkova O, Fazekas A, Macias VM, Bier E, James AA. 2015. Highly efficient Cas9-mediated gene drive for population modification of the malaria vector mosquito *Anopheles stephensi*. Proceedings of the National Academy of Sciences 112(49): E6736–E6743.

Gardiner DM, Kazan K. 2018. Selection is required for efficient Cas9-mediated genome editing In *Fusarium graminearum*. Fungal Biology 122(2-3): 131–137.

Gardiner DM, Stephens AE, Munn AL, Manners JM. 2013. An ABC pleiotropic drug resistance transporter of *Fusarium graminearum* with a role in crown and root diseases of wheat. FEMS Microbiology Letters 348(1): 36–45.

Gardiner DM, Stiller J, Kazan K. 2014. Genome sequence of *Fusarium graminearum* isolate CS3005. Genome Announcements 2(2): e00227–00214.

Gietz RD, Schiestl RH. 2007. High-efficiency yeast transformation using the LiAc/SS carrier DNA/PEG method. Nature Protocols 2(1): 31–34.

Girardin L, Calvez V, Débarre F. 2019. Catch Me If You Can: A Spatial Model for a Brake-Driven Gene Drive Reversal. Bulletin of Mathematical Biology.

Gladyshev E 2017. Repeat-induced point mutation and other genome defense mechanisms in fungi.In Heitman J, Stukenbrock EH. Microbial Spectrum. Washington DC: American Society of Microbiology.

Goswami RS, Kistler HC. 2004. Heading for disaster: *Fusarium graminearum* on cereal crops. Molecular Plant Pathology 5(6): 515–525.

Grognet P, Lalucque H, Malagnac F, Silar P. 2014. Genes that bias mendelian segregation. PLoS Genetics 10(5): e1004387.

Grunwald HA, Gantz VM, Poplawski G, Xu X-RS, Bier E, Cooper KL. 2019. Super-Mendelian inheritance mediated by CRISPR–Cas9 in the female mouse germline. Nature 566: 105–109.

Hammond A, Galizi R, Kyrou K, Simoni A, Siniscalchi C, Katsanos D, Gribble M, Baker D, Marois E, Russell S, et al. 2016. A CRISPR-Cas9 gene drive system targeting female reproduction in the malaria mosquito vector *Anopheles gambiae*. Nat Biotech 34(1): 78–83.

Jones MS, Delborne JA, Elsensohn J, Mitchell PD, Brown ZS. 2019. Does the U.S. public support using gene drives in agriculture? And what do they want to know? Science Advances 5(9): eaau8462.

Kazan K, Gardiner DM, Manners JM. 2012. On the trail of a cereal killer: recent advances In *Fusarium graminearum* pathogenomics and host resistance. Molecular Plant Pathology 13(4): 399–413.

King R, Urban M, Hammond-Kosack MCU, Hassani-Pak K, Hammond-Kosack KE. 2015. The completed genome sequence of the pathogenic ascomycete fungus *Fusarium graminearum*. BMC Genomics 16(1): 1–21.

Laurent B, Palaiokostas C, Spataro C, Moinard M, Zehraoui E, Houston RD, Foulongne-Oriol M. 2018. High-resolution mapping of the recombination landscape of the phytopathogen *Fusarium graminearum* suggests two-speed genome evolution. Molecular Plant Pathology 19(2): 341–354.

Lindholm AK, Dyer KA, Firman RC, Fishman L, Forstmeier W, Holman L, Johannesson H, Knief U, Kokko H, Larracuente AM, et al. 2016. The ecology and evolutionary dynamics of meiotic drive. Trends in Ecology & Evolution 31(4): 315–326.

Lofgren LA, LeBlanc NR, Certano AK, Nachtigall J, LaBine KM, Riddle J, Broz K, Dong Y, Bethan B, Kafer CW, et al. 2018. *Fusarium graminearum*: pathogen or endophyte of North American grasses? New Phytologist 217(3): 1203–1212.

Malz S, Grell MN, Thrane C, Maier FJ, Rosager P, Felk A, Albertsen KS, Salomon S, Bohn L, Schäfer W, et al. 2005. Identification of a gene cluster responsible for the biosynthesis of aurofusarin in the *Fusarium graminearum* species complex. Fungal Genetics and Biology 42(5): 420–433.

Nuckolls NL, Bravo Núñez MA, Eickbush MT, Young JM, Lange JJ, Yu JS, Smith GR, Jaspersen SL, Malik HS, Zanders SE. 2017. *wtf* genes are prolific dual poison-antidote meiotic drivers. eLife 6: e26033.

Orton ES, Deller S, Brown JKM. 2011. *Mycosphaerella graminicola*: from genomics to disease control. Molecular Plant Pathology 12(5): 413–424.

Pomraning KR, Connolly LR, Whalen JP, Smith KM, Freitag M 2013. Repeat-induced point mutation, DNA methylation and heterochromatin in Gibberella zeae (anamorph: Fusarium graminearum). In: Brown DW, Proctor RH eds. Fusarium: genomics, molecular and cellular biology. Norfolk, United Kingdom: Caister Academic Press, 93–109.

Prieto M, Wedin M. 2013. Dating the diversification of the major lineages of Ascomycota (fungi). PLoS ONE 8(6): e65576.

Rouxel T, Balesdent MH. 2005. The stem canker (blackleg) fungus, *Leptosphaeria maculans*, enters the genomic era. Molecular Plant Pathology 6(3): 225–241.

Svedberg J, Hosseini S, Chen J, Vogan AA, Mozgova I, Hennig L, Manitchotpisit P, Abusharekh A, Hammond TM, Lascoux M, et al. 2018. Convergent evolution of complex genomic rearrangements in two fungal meiotic drive elements. Nature Communications 9(1): 4242.

Trail F. 2009. For blighted waves of grain: *Fusarium graminearum* in the postgenomics era. Plant Physiology 149(1): 103–110.

Vogan AA, Ament-Velásquez SL, Granger-Farbos A, Svedberg J, Bastiaans E, Debets AJM, Coustou V, Yvanne H, Clavé C, Saupe SJ, et al. 2019. Combinations of *Spok* genes create multiple meiotic drivers In Podospora. eLife 8: e46454.

Ward TJ, Clear RM, Rooney AP, O’Donnell K, Gaba D, Patrick S, Starkey DE, Gilbert J, Geiser DM, Nowicki TW. 2008. An adaptive evolutionary shift In *Fusarium* head blight pathogen populations is driving the rapid spread of more toxigenic *Fusarium graminearum* in North America. Fungal Genetics and Biology 45(4): 473–484.

Wedell N, Price TAR, Lindholm AK. 2019. Gene drive: progress and prospects. Proceedings of the Royal Society B: Biological Sciences 286(1917): 20192709.

Wortman JR, Gilsenan JM, Joardar V, Deegan J, Clutterbuck J, Andersen MR, Archer D, Bencina M, Braus G, Coutinho P, et al. 2009. The 2008 update of the *Aspergillus nidulans* genome annotation: A community effort. Fungal Genetics and Biology 46 (1, Supplement): S2-S13.

